# Brain inflammation and cognitive decline induced by spinal cord injury can be reversed by spinal cord cell transplants

**DOI:** 10.1101/2024.09.20.614106

**Authors:** Quentin Delarue, Amandine Robac, Célia Duclos, Baptiste Pileyre, Clémence Raimond, Pauline Neveu, Gaëtan Riou, Nicolas Guérout

## Abstract

Spinal cord injuries (SCIs) affect between 250,000 and 500,000 people worldwide each year, most commonly due to road accidents or falls. These injuries result in permanent disabilities, the severity and impact of which are directly related to the extent and location of the injury. Recent studies have also shown that SCIs can lead to cognitive disorders due to inflammation in the brain.

From a therapeutic perspective, numerous treatments have been explored, including cell therapy. It has been established that a common mechanism across various cellular transplant models is the modulation of inflammation at the injury site. However, it remains unclear whether the immunomodulatory effects observed in the spinal cord also extend to the brain.

To test this hypothesis, we induced SCI in wild-type mice and treated them with transplants of differentiated cells, specifically olfactory ensheathing cells, or stem cells, such as mesenchymal stem cells. Our results demonstrate that both types of transplants can reverse cognitive disorders induced by SCI. Additionally, we found that these cellular transplants modulate brain inflammation and increase neuronal density in the hippocampus.

To our knowledge, this is the first study to show that cells transplanted into the spinal cord can modulate the inflammatory response in the brain, thereby reversing the negative effects of injury on brain function following SCI. These findings underscore the complex interactions between the brain and spinal cord under both physiological and pathological conditions.

## INTRODUCTION

Spinal cord injuries (SCIs) are most commonly caused by trauma from road accidents or falls. They predominantly affect men, with the highest incidence among young adults and those over 60 (Golestani et al., 2022; Liu et al., 2023). Depending on their severity and location, SCIs result in the loss of motor, sensory, and autonomic functions, and currently, there is no cure.

The pathophysiology of SCIs has been studied for many years and is well documented (Anjum et al., 2020). More recently, research has focused on the effects of SCIs on the brain, showing that they also cause inflammation in brain parenchyma. This inflammation is responsible for a reduction in hippocampal neurogenesis, leading to cognitive deficits in human and mice (Craig et al., 2017; Li et al., 2020; Maldonado-Bouchard et al., 2016). Therapeutic approaches most frequently studied to repair damaged spinal cords include cell transplantation (Assinck et al., 2017) and, more recently, electrical (Wagner et al., 2018) and magnetic neuromodulation techniques (Chalfouh et al., 2020; Robac et al., 2021). Historically, various cell types have been investigated in the context of SCIs, including differentiated cells like bulbar olfactory ensheathing cells (bOECs) and stem cells such as mesenchymal stem cells (MSCs). For the past thirty years, bOECs have been studied in preclinical animal studies and clinical human trials in the context of SCIs (Gómez et al., 2018). It has been shown that these cells can modulate spinal cord scar, promote the regrowth or survival of motor fibers, and enable functional recovery without integrating into the spinal tissue (Delarue et al., 2021; Wang et al., 2022). Interestingly, a growing body of research indicates that the primary role of bOECs after transplantation is to exert an immunomodulatory effect on microglia and macrophages present at the lesion site (Khankan et al., 2016; Zhang et al., 2021). Regarding MSCs, these non-neural cells have also shown great interest in tissue regeneration after SCI in animal models and humans. They stimulate spinal cord tissue regeneration, axonal regrowth (Chen et al., 2022; Tien et al., 2019) and like bOECs, provide an immunomodulatory effect (Han et al., 2015; Nakajima et al., 2012).

Based on this knowledge, we asked ourselves: just as inflammation in the spinal cord can lead to disruptions in the brain following injury, can the immunomodulatory effects of bOECs or MSCs transplanted into the spinal cord after SCI induce beneficial effects on brain tissue and help restore in the recovery of cognitive performance?

Therefore, our study represents the first investigation into the impact of spinal cord transplantation of bOECs or MSCs following SCI on brain inflammation and cognitive functions.

## MATERIALS AND METHODS

### Animal care and use statement

The animal protocol was designed to minimize pain and discomfort for animals. All experimental procedures complied with the European Community guiding principles on the care and use of animals (86/609/CEE; Official Journal of the European Communities No. L358; December 18, 1986), the French Decree No. 97/748 of October 19, 1987 (Journal Officiel de la République Française; October 20, 1987), and the recommendations of the Cenomexa Ethics Committee (#26905).

### Animals

Mice of both sexes were group-housed (two to five mice per cage, with genders separated) in secure conventional rodent facilities, maintained on a 12-hour light/dark cycle with constant access to food and water. A total of 81 wild-type C57BL/6 mice were included in this study. All experiments were conducted on adult mice aged 8–12 weeks, with an average weight of 20 g for females and 25 g for males. Experimental groups were formed with an equal sex ratio (50:50). Our study consisted of four main experimental groups:

Non-injured control group: mice without SCI used to establish baselines in histological and functional experiments (Fig. 1a and 2a).

**Figure 1:**
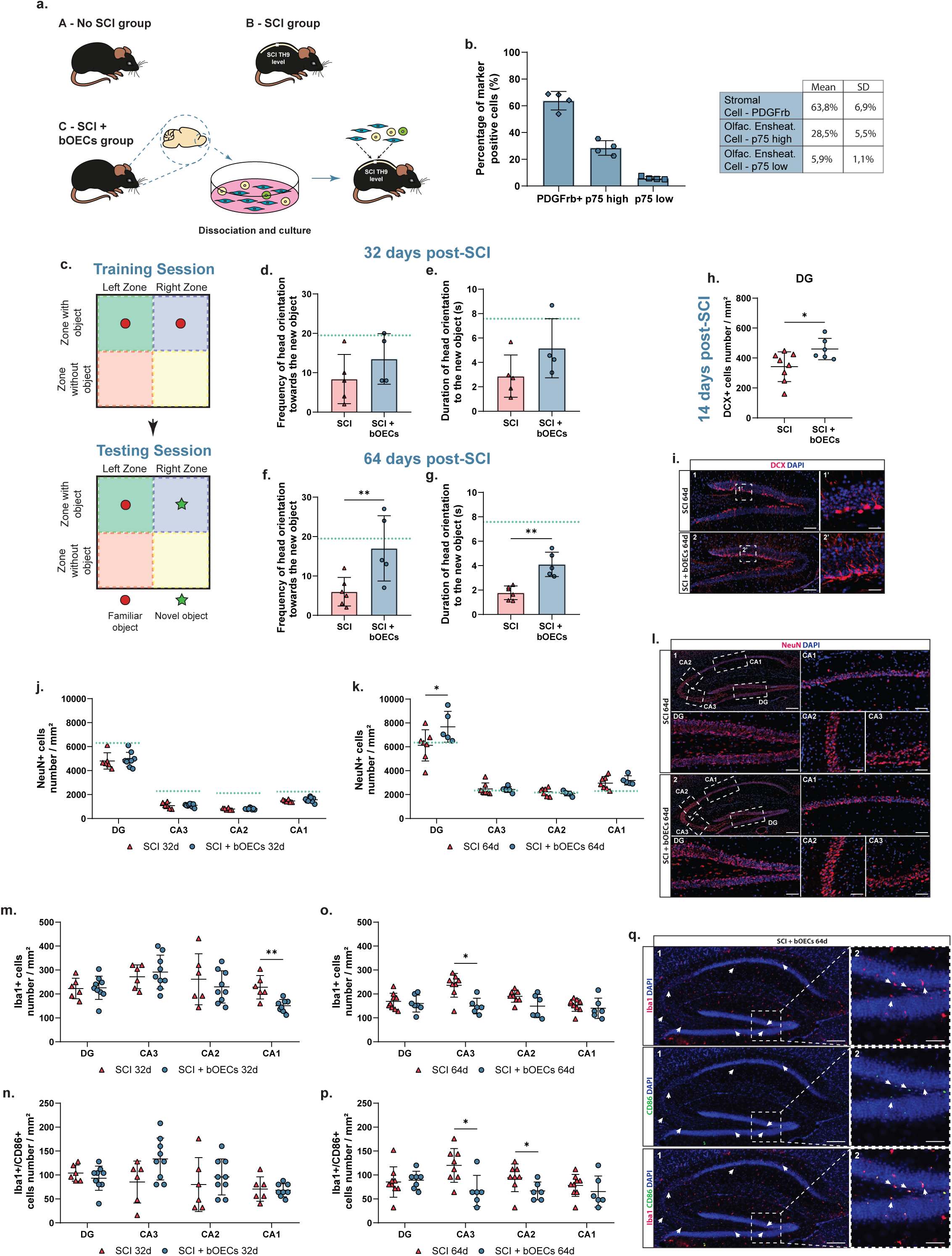
Transplantation of bOECs reduces hippocampal inflammation and corrects the memory deficit induced by SCI. **a.** Schematic representation of the three experimental groups: A -No SCI group, B - SCI group, and C - SCI + bOECs group. **b.** Cytometric characterization of bOECs primary culture. Anti-PDGFrβ and anti-p75 antibodies were used to identify stromal cells and OECs, respectively. **c-g.** NOR experiments were conducted 32 and 64 days after SCI. **c.** Schematic representation of the NOR experiment. After a training session with two identical objects, one object was replaced by a novel object different from the first. During the testing session, the frequency and duration of head orientation towards the novel object were measured. **d and f.** Frequency of head orientation towards the novel object at 32 (**d**) and 64 (**f**) days post-SCI. **e and g.** Duration of head orientation towards the novel object at 32 (**e**) and 64 (**g**) days post-SCI. **h-q.** Immunohistology experiments. **h and i.** Quantification of immature neurons (DCX+) in the DG 14 days post-SCI. **h.** Quantification of DCX+ neuron density in the DG 14 days post-SCI. **i.** Representative images of sagittal DG sections from SCI (**i1 and i1’**) and SCI + bOECs (**i2 and i2’**) groups 14 days post-SCI, stained with anti-DCX antibody. **i1 and i2:** Scale bar = 100 µm; **i1’ and i2’**: Scale bar = 50 µm. **j and k.** Quantification of mature neuron (NeuN+) density in the DG, CA3, CA2, and CA1 at 32 (**j**) and 64 (**k**) days post-SCI. **l.** Representative images of hippocampal sagittal sections from the SCI (**l1**) and SCI + bOECs (**l2**) groups, stained with anti-NeuN antibody. **l1 and l2**: Scale bar = 100 µm; **DG, CA1, CA2, and CA3**: Scale bar = 50 µm. **m-q.** Analysis of microglial cell reactivity in the hippocampus 32 and 64 days post-SCI. **m and o.** Quantification of microglial cell density (Iba1+) in the hippocampus at 32 (**m**) and 64 (**o**) days post-SCI. **n and p.** Quantification of microglial cell reactivity (Iba1+/CD86+) in the hippocampus at 32 (**n**) and 64 (**p**) days post-SCI. **q.** Representative images of hippocampal sagittal sections from SCI + bOECs, stained with anti-Iba1 and anti-CD86 antibodies. Arrows indicate Iba1+/CD86+ microglial cells. **q1**: Scale bar = 100 µm; **q2**: Scale bar = 50 µm. Quantifications are expressed as mean ± SD (**b, d, e, f, g, h, j, k, m, and o**). N = 5 (**b**), 5-6 (**d-g**), 6-8 (**h**), and 6-9 (**j-o**) animals per group. Statistical evaluations were based on Mann-Whitney tests. * = P < 0.05; ** = P < 0.01.

**Figure 2:**
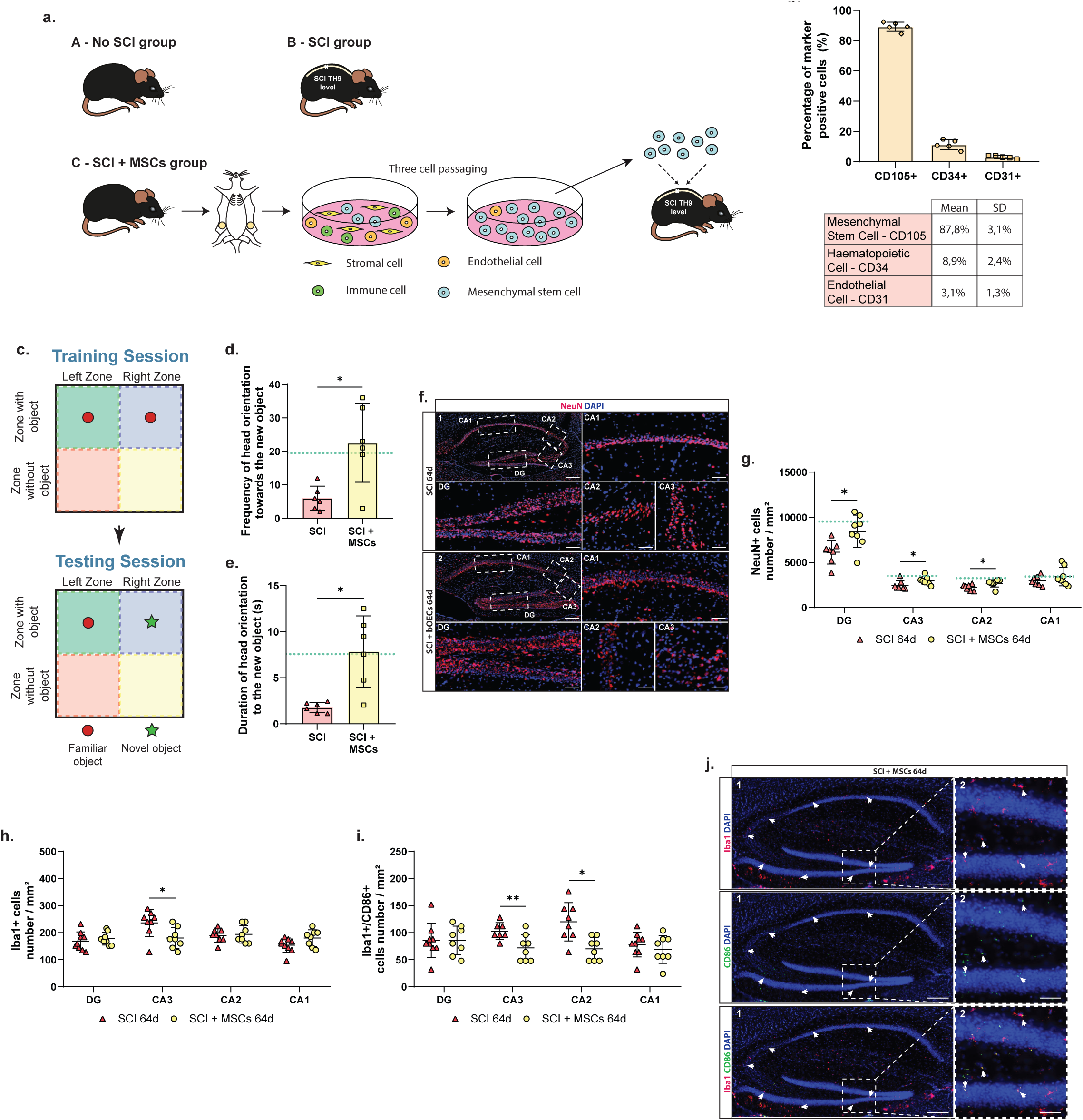
Transplantation of MSCs also corrects the memory deficit induced by SCI by acting on the same structures and cells as the transplantation of bOECs. **a.** Schematic representation of the three experimental groups: A - No SCI group, B - SCI group, and C - SCI + MSCs group. **b.** Cytometric characterization of MSCs purified culture. Anti-CD105, anti-CD34 and anti-CD31 antibodies were used to identify MSCs, hematopoietic cells and endothelial cells, respectively. **c-e.** NOR experiments were conducted 64 days after SCI. **c.** Schematic representation of the NOR experiment. Like for bOECs transplantation experiment after a training session with two identical objects, one object was replaced by a novel object different from the first. During the testing session, the frequency and duration of head orientation towards the novel object were measured. **d.** Frequency of head orientation towards the novel object at 64 days post-SCI. **e.** Duration of head orientation towards the novel object at 64 days post-SCI. **f-j.** Immunohistology experiments. **f.** Representative images of hippocampal sagittal sections from the SCI (**f1**) and SCI + MSCs (**f2**) groups, stained with anti-NeuN antibody. **f1 and f2**: Scale bar = 100 µm; **DG, CA1, CA2, and CA3**: Scale bar = 50 µm. **g.** Quantification of mature neuron (NeuN+) density in the DG, CA3, CA2, and CA1 at 64 days post-SCI. **h-j.** Analysis of microglial cell reactivity in the hippocampus 64 days post-SCI. **h.** Quantification of microglial cell density (Iba1+) in the hippocampus at 64 days post-SCI. **i.** Quantification of microglial cell reactivity (Iba1+/CD86+) in the hippocampus at 64 days post-SCI. **j.** Representative images of hippocampal sagittal sections from SCI + MSCs groups, stained with anti-Iba1 and anti-CD86 antibodies. Arrows indicate Iba1+/CD86+ microglial cells. **j1**: Scale bar = 100 µm; **j2**: Scale bar = 50 µm. Quantifications are expressed as mean ± SD (**b, d, e, g, h, and i**). N = 5 (**b**), 6 (**d and e**) and 8-9 (**h and i**) animals per group. Statistical evaluations were based on Mann-Whitney tests. * = P < 0.05; ** = P < 0.01.

SCI control group: mice underwent SCI without transplantation (Fig. 1a and 2a).

SCI + bOECs group: mice underwent SCI and received primary culture of bOECs (Fig. 1a).

SCI + MSCs group: mice underwent SCI and received MSCs (Fig. 2a).

### Primary olfactory bulb culture and cell transplant preparation

Olfactory bulb (OB) primary cultures were prepared as previously described (Delarue et al., 2019; Guérout et al., 2010). Briefly, mice were anesthetized with 2% isoflurane (Iso-Vet, Osalia, Paris, France) and then euthanized by decapitation. OBs were extracted from the brain and placed in cold PBS. The tissue was incubated in 0.25% trypsin (2.5% Trypsin 10X, 15090-046, Thermo Fisher) and mechanically dissociated. Once dissociated, cells were cultured in T25 flasks with 5 mL of DF-10S medium (DMEM + GlutaMAX, 0.5% Penicillin/Streptomycin, and 10% heat-inactivated fetal bovine serum). Three weeks after plating, cultures were trypsinized and cells were counted to prepare a cell suspension at 25,000 cells/µL in DF-10S for transplantation.

### Mesenchymal stem cell culture and cell transplant preparation

MSCs cultures were prepared from adipose tissue. Briefly, mice were anesthetized with 2% isoflurane (Iso-Vet, Osalia, Paris, France) and then euthanized by decapitation. Adipose tissue from the abdomen was isolated and placed in cold PBS. The tissue was minced with a syringe and dissociated using NB5 collagenase (Nordmak, N0002778) on an orbital shaker for 1 hour. Once dissociated, cells were cultured in T75 flasks with 15 mL of medium (DMEM + GlutaMAX, 1% Penicillin/Streptomycin, and 20% heat-inactivated fetal bovine serum). One week after plating, cultures were trypsinized and replated in T75 flasks. Cultures were passaged three times to obtain purified MSCs. After three passages, cells were counted, and a cell suspension was prepared at 50,000 cells/µL in culture medium for transplantation.

### Flow cytometry and bOECs characterization

Cells were characterized by flow cytometry three weeks after plating. Primary bOECs were adjusted to a density of 2 × 10□ cells/mL in PBS with 0.5% bovine serum albumin (BSA) solution. TruStain FcX™ PLUS (BioLegend) was used to block non-specific binding. Cell populations were identified using primary antibodies: anti-p75 nerve growth factor receptor (p75, Abcam, ab8874, RRID) and rat anti-platelet-derived growth factor β (PDGFrβ, Abcam, ab91066, RRID). OECs were identified as p75-positive and PDGFrβ-negative cells, while stromal cells were identified as PDGFrβ-positive and p75-negative cells. p75 and PDGFrβ primary antibodies were revealed with anti-rabbit phycoerythrin (PE, poly4064, BioLegend, 406408, RRID) and anti-rat Alexa Fluor 488 (AF488, MRG2b-85, BioLegend, RRID) fluorochrome-conjugated secondary antibodies, respectively. Data were analyzed using FlowJo software (version 10.3; FlowJo LLC).

### Flow cytometry and MSCs characterization

Cells were characterized by flow cytometry after three passages. MSCs were adjusted to a density of 2 × 10□ cells/mL in PBS with 0.5% BSA solution. TruStain FcX™ PLUS (BioLegend) was used to block non-specific binding. MSCs were characterized as CD105-positive, CD31-negative, and CD34-negative cells. In contrast, endothelial cells were defined as CD105-negative, CD31-positive, and CD34-negative, while hematopoietic cells were identified as CD105-negative, CD31-negative, and CD34-positive. Cells were labeled with the following antibodies: anti-endoglin conjugated with peridinin-chlorophyll-protein complex-cyanine5.5 (CD105-PerCP-Cy5.5, Sony, 1202080), anti-PECAM1 conjugated with phycoerythrin (CD31-PE, Sony, 1112040), and anti-mucosialin conjugated with allophycocyanin (CD34-APC, Sony, 1243060). Data were analyzed using FlowJo software (version 10.3; FlowJo LLC).

### Surgical procedure and cell transplantation

SCI was performed at the T9-T10 level as previously described (Delarue et al., 2019). After a dorsal incision and laminectomy at the T7 vertebra, an incomplete transection of the spinal cord was performed to preserve the lower part of the ventral horns while completely injuring the central part and the dorsal horns. The SCI was made using a 25-gauge needle, and immediately following the lesion, bOECs or MSCs were transplanted. Both bOECs and MSCs were transplanted using a micromanipulator arm fixed to a stereotaxic frame (World Precision Instruments). Cells were injected with a 1 mm sterile glass capillary needle attached to the micromanipulator. Injections were delivered at a 1 mm depth, 1.5 mm from the midline, and 5 mm rostrally and caudally from the lesion site. Two injections of 2 µL each, containing 25,000 cells/µL of bOECs or 50,000 cells/µL of MSCs, were administered. Each injection was carefully delivered over 1 minute. After surgery, back muscles and skin were sutured, and mice were monitored daily. None of the animals showed signs of skin lesions, infection, or autophagy throughout the study.

### Novel object recognition (NOR)

To assess non-spatial hippocampal-mediated memory, the NOR test was performed. The primary aim of this test is to evaluate the mouse’s ability to recognize a novel object. Briefly, the day before the test, mice were individually placed twice, at the beginning and end of the day, in a 35 × 35 cm open field for 5 minutes each time for habituation. The next day, mice were placed in the open field with two identical objects for 10 minutes (training session). After 2 hours, object recognition was assessed by replacing one of the familiar objects with a novel object that differed in color, shape, size, and texture. Mice were then individually placed in the same open field for 10 minutes, and the time spent exploring the familiar and novel objects was measured using a camera positioned above the field (testing session) (Fig 1c and 2c). Since mice have an inherent preference for exploring novel objects, a preference for the novel object indicates intact memory for the familiar object. Experiments were conducted 32 and 64 days after SCI.

### Tissue preparation and sectioning

Thirty minutes before euthanasia, mice received analgesic treatment to minimize pain (Buprenorphine 0.17 mg/kg). Following this, mice were deeply anesthetized with 5% isoflurane and euthanized via intraperitoneal injection of Euthoxin (100 mg/kg). Transcardial perfusion with cold PBS was followed by ice-cold 4% paraformaldehyde (PFA, Sigma Aldrich, 252549) in PBS. The dissected spinal cord and brain were then post-fixed in 4% PFA overnight and cryoprotected in 30% sucrose (Life Technologies, Carlsbad, CA) for at least 48 hours. After embedding in Tissue-Tek OCT compound (Sakura, Tokyo, Japan), brains were cut sagittally into 30 µm thick sections. Sections were collected on slides and stored at -20°C until further use.

### Immunohistochemistry

Identification of different cells and structures in the brain was performed using primary antibodies diluted in a PBS-blocking solution containing 10% normal donkey serum (Jackson ImmunoResearch, Cambridge, UK) and 0.3% Triton-X100 (Sigma-Aldrich) to block non-specific binding. Antibody incubation was conducted overnight at room temperature in a humidified chamber. The following primary antibodies were used: rabbit anti-ionized calcium-binding adapter molecule 1 (Iba1, Wako, 019-19741, Osaka, Japan, RRID), rat anti-B-lymphocyte activation antigen B7-2 (CD86, Invitrogen, MA1-10299, RRID:AB_11152324), mouse anti-RNA binding Fox-1 homolog 3 (NeuN, Sigma, HPA030790, RRID) and goat anti-doublecortin (DCX, Santa Cruz, SC-8066, RRID).

After washing, antibody staining was revealed using species-specific fluorescence-conjugated secondary antibodies (Jackson ImmunoResearch). Sections were counterstained with 4′,6-diamidino-2-phenylindole (DAPI; 1 μg/mL; Sigma-Aldrich) and coverslipped with Vectashield mounting media (Vector Labs, Burlingame, UK).

### Image acquisition analysis

Representative images for quantification of immunohistochemically stained areas in sagittal sections were captured using the Zeiss Apotome2 and Leica THUNDER Imager Tissue 3D microscope setups. For brain analysis, hippocampus images were obtained using a 20x objective lens. Image processing and assembly were performed using ImageJ software.

### Quantification of immunohistochemically stained areas

The depth and number of brain sections analyzed were determined based on the depth and thickness of the granular cell layer in the OB. In the hippocampal sections, Iba1+, CD86+, DCX+, and NeuN+ cells were counted using standardized rectangular regions for each hippocampal area. For the dentate gyrus (DG) and cornu Ammonis (CA) 1, cells were counted within rectangles of 500 µm × 250 µm. For the CA2 and CA3 regions, cells were counted within rectangles of 250 µm × 250 µm.

### Statistical analysis

All data are presented as mean ± standard deviation (SD). Statistical analyses were performed using GraphPad Prism software, version 8.0.1.244 for Windows (GraphPad Software). The comparison of means was conducted using the two-way Mann-Whitney test. Detailed statistical analyses for each assay are provided in the figure legends. A p-value of < 0.05 was considered statistically significant.

## RESULTS

### Transplantation of bOECs into the spinal cord improves cognitive behavior in mice after SCI

As previously described, the emergence of the cerebral inflammatory response after SCI is linked to cognitive deficits causing memory impairment in mice (Li et al., 2020). In a previous study, we demonstrated that spinal cord transplantation of bOECs reduced the inflammatory response in the spinal cord parenchyma (Delarue et al., 2021). Based on these results, we have investigated the effects of bOECs transplantation in spinal cord after SCI on cognitive behavior, hippocampal neurogenesis and inflammation. First, we performed flow cytometry cellular characterization of the primary culture of bOECs obtained from the OB of mice. Using PDGFrβ and P75 antibodies to identify stromal cells and OECs respectively, we found that our primary cultures consist of 63.8% ± 6,9% stromal cells and 31.4% ± 6,6 OECs. OECs are divided into two subpopulations: p75 high and p75 low OECs, representing 28.5% ± 5,5% and 5.9% ± 1,1%, respectively (Fig. 1b). These findings are consistent with previous studies on murine models, which have shown that primary cultures of bOECs consist of approximately 60% to 70% stromal cells and 30% to 40% OECs (Gómez et al., 2018).

Then, SCI and bOECs transplantation were performed at the same time, and longitudinal NOR experiments were conducted 32 and 64 days post-SCI (Fig. 1c-g). This cognitive test, based on the ability to recognize the presence of a new object in their environment, measures the memory capacity of mice. As previously described, NOR results indicate that SCI induces significant memory capacity decline, evidenced by the reduced frequency and duration of head orientation towards the novel object in the SCI group compared to the non-injured control group (dashed lines) 32 and 64 days post-SCI (Fig. 1d-g). Notably, our results demonstrate that bOECs transplantation ameliorates memory capacity decline when compared to the SCI group. Indeed, 64 days after SCI, the frequency and duration of head orientation towards the new object are significantly increased in the bOECs group (Fig. 1f and g), but not at 32 days (Fig. 1d and e).

To investigate the mechanisms underlying memory improvement in transplanted mice, we collected brains from mice that had performed the NOR test at 64 days post-SCI. Additionally, in order to study the effects of bOECs transplantation on brain at earlier stages, we transplanted bOECs into other groups of mice after SCI and collected their brains either 14 or 32 days after SCI.

Firstly, we investigated hippocampal neurogenesis. Indeed, in mice, adult hippocampal neurogenesis plays a crucial role in maintaining learning and memory mechanisms (Berdugo-Vega et al., 2020). Using immunohistochemistry, we measured the number of immature (DCX+) and mature (NeuN+) neurons in the four hippocampal regions: DG, CA1, CA2, and CA3. These analyses reveal that bOECs transplantation increases the number of DCX+ cells in the hippocampus 14 days after SCI (Fig. 1h and i). Additionally, bOECs transplantation increases the number of NeuN+ cells specifically in the DG 64 days after SCI (Fig. 1k and l), with no effect observed at 32 days post-SCI (Fig. 1j). These interesting results highlight a potential increase in neurogenesis during the first two weeks following the injury, leading to the generation of mature neurons in the transplanted mice.

Synaptic plasticity is a crucial factor in maintaining cognitive functions. The inflammatory response negatively affects the memory capacity of mice by reducing synaptic plasticity (Prieto et al., 2019). We therefore explored the effect of bOECs transplantation on microglial activation in the hippocampus. To do this, we used Iba1 and CD86 antibodies to quantify microglial cell reactivity through immunohistochemistry at 32 and 64 days post-SCI. The number of Iba1+ cells is decreased in CA1 at 32 days (Fig. 1m) and in CA3 at 64 days (Fig. 1o and q) post-SCI in bOECs transplanted mice compared to non-transplanted mice. Additionally, bOECs transplantation affects microglial activation 64 days after SCI by reducing the number of Iba1+/CD86+ activated microglial cells in CA2 and CA3 (Fig. 1p and q), but without effect 32 days post SCI (Fig. 1n).

Altogether, our results demonstrate that bOECs transplantation can restore non-spatial hippocampal-mediated memory decline associated with SCI, potentially by increasing hippocampal neurogenesis in the DG and promoting synaptic plasticity through the reduction of microglial activation in the CA regions.

### Improvement of cognitive functions after SCI is also facilitated by non-neural cells transplantation

To confirm whether the improvement in memory capacity is enabled by the transplantation of other cells, particularly those of non-neural origin, we conducted the same experiments and analyses on mice that received MSCs derived from adipose tissue following SCI. To do this, after cell culture characterization we performed the NOR experiment, quantified the number of neurons, and assessed hippocampal microglial reactivity 64 days after SCI.

MSCs were obtained from inguinal adipose tissue. After three passages, cells were characterized by flow cytometry using CD105, CD31, and CD34 antibodies to identify MSCs, endothelial cells, and hematopoietic cells, respectively. We identified 87.8% MSCs, 3.1% endothelial cells, and 8.9% hematopoietic cells (Fig. 2b), which is consistent with previous studies (Luna et al., 2014).

We then transplanted MSCs into mice following SCI and conducted the NOR experiment. Interestingly, similar to bOECs transplantation experiments, MSCs transplantation at 64 days post-SCI compensates for the memory deficit induced by SCI (Fig. 2c-e). Although we did not quantify the number of immature DCX+ neurons 14 days after SCI, it is possible that the improvement in cognitive performance is also linked to the induction of hippocampal neurogenesis in mice transplanted with MSCs. Indeed, the number of mature NeuN+ neurons is increased in the DG 64 days post-SCI and transplantation. Unlike mice transplanted with bOECs, we also observed an increase in the number of mature neurons in CA2 and CA3 (Fig. 2f and g).

Regarding the effect of MSCs transplantation on hippocampal inflammation, we also observed a reduction in the number of microglial cells in CA3, identified by Iba1 (Fig. 2h and j), as well as a decrease in the number of Iba1+/CD86+ cells in CA2 and CA3, 64 days after SCI, in mice that received MSCs compared to non-transplanted mice (Fig. 2i and j). This indicates a reduction in hippocampal microglial reactivity and potentially a correction of the synaptic plasticity inhibition induced by SCI.

These results highlight the ability of MSCs transplantation into the spinal cord after SCI, similar to bOECs transplantation, to improve memory deficits induced by SCI. These effects appear to be associated with an increase in hippocampal neurogenesis as well as a reduction in microglial reactivity.

## DISCUSSION

### Therapeutic effects of bOECs transplantation after SCI on brain reactivity and disturbances of the cognitive functions

The purpose of our study was to investigate the effect of cell transplantation, bOECs and MSCs, used as a treatment after SCI on brain reactivity and cognitive functions. As described above, SCI induces microglia brain activation particularly in hippocampus (Li et al., 2020). Studies of anxiety, depression and cognitive performances of patients suffering SCI, show that it induces brain perturbations lasting for months after the injury. The importance of these perturbations is correlated with the severity of SCI, conditioning the cognitive deficits after the traumatism. Patients with high cognitive deficits, induced by SCI, have the worst cognitive recovery (Craig et al., 2017; Maldonado-Bouchard et al., 2016). These perturbations are related to microglia activation, causing a reduction of neuronal density which is linked to a decrease of neurogenesis and an increase of neuronal death. In rodents, specific test as NOR, allow to investigate brain reactivity and cognitive performances (Li et al., 2020). Based on these different studies, we carried out cognitive test after SCI with bOECs transplantation, MSCs transplantation or without treatment. Our findings show for the first time that cellular therapy applied at spinal level modulates the brain reactivity induced by SCI. Indeed, bOECs and MSCs transplantation enhances memory performance 64 days after SCI, objectified by NOR test (Fig. 1f-g and 2d-e). To elucidate how bOECs transplantation can enhance memory performance, we have performed histological analyses of hippocampus to quantify neurogenesis at early stage. Neurogenesis in DG is perturbated by SCI and is enhanced by bOECs transplantation 14 days after SCI (Fig. 1h and i). Based on the time required for a neuron to mature, we hypothesize that these new immature DCX+ neurons contribute to increase the number of mature NeuN+ neurons observed 64 days after SCI, as we did not observe any difference in neuronal density in the DG 32 days after SCI (Fig. 1j and k). This hypothesis is further supported by the fact that neurogenesis allows the generation of new neurons exclusively in the DG (Llorens-Martín et al., 2015; Zhao et al., 2007).

To confirm the beneficial effects of bOECs transplantation on the brains of mice who underwent SCI, we analyzed the hippocampus of mice transplanted with MSCs 64 days after SCI. Interestingly, we also observed an increase in the density of mature neurons NeuN+ in the DG, as well as in the CA2 and CA3 regions (Fig. 2g). This could possibly be explained by an increase in neuronal survival in the MSCs-transplanted mice.

To investigate other mechanisms which can modulate cognitive functions, we have assessed the hippocampal inflammatory response. Microglial cells participate to hippocampal homeostasis by modulating neurogenesis, synaptogenesis and neuronal death (Li et al., 2020). That is why, we have then, investigated microglial reactivity into the hippocampus. Our results, show that bOECs transplantation reduces microglial activation in CA1 area at the beginning of the chronic stage (Fig. 1m). CA1 area is the starting point for neuronal projections leaving the hippocampus, it is important for memory transmission in other brain areas. Thus, the disruption of CA1 area brings memory deficits, particularly for non-spatial learning as object recognition, studied with NOR test (Basu and Siegelbaum, 2015). However, we did not find significative differences in NOR test results at 32 days post-SCI. On the other hand, at chronic stage, bOECs and MSCs transplantation has no effect on microglial reactivity in CA1 but reduces it in CA3 (Fig. 1o-q). CA3 area is involved in episodic memory, encoding spatial representation and the coordination of hippocampal areas (Cherubini and Miles, 2015). Moreover, CA3-CA1 synaptic connections control long-term potentiation (LTP) generation in CA1 neurons. LTP generation in CA1 is correlated with learning and memory (Whitlock et al., 2006).

Altogether, our results show that bOECs and MSCs transplantation in spinal cord after injury induces therapeutic effects on memory deficits caused by SCI. This effect is potentially associated with improvement of neurogenesis at early stage based on the increase of the number of DCX+ neurons. Moreover, bOECs and MSCs transplantation enhances the number of mature neurons (NeuN+) at chronic stage (Fig. 1k and 2g). These results regarding mature neurons can be interpreted as an improvement of neuronal integration of the newborn neurons (DCX+) observed 14 days after SCI. Finally, our results show also that bOECs and MSCs transplantation decreases microglial cells reactivity in CA1 and CA3, favorizing the synaptic plasticity and learning.

This study does not allow for a precise description of the mechanisms involved, but several hypotheses can be proposed. One plausible explanation is the involvement of inflammatory cytokines propagation to the brain. Cytokines, chemokines, and reactive oxygen species (ROS) are present in the cerebrospinal fluid (CSF) and blood after SCI, and they can increase the permeability of the blood-brain barrier (BBB), facilitating the infiltration of activated immune cells into the brain (Anthony et al., 2014; Capirossi et al., 2020; McCandless et al., 2008). Endothelial cells, which compose the BBB, express transporters for cytokines and chemokines (Banks et al., 1994; Osburg et al., 2002). Additionally, lymphocytes can transiently enter the brain through various routes, including the non-windowed capillaries of the choroid plexus. As lymphocytes constitute 50% of immune cells in the CSF, they can be activated by cytokines in the CSF and, in turn, activate cerebral microglial cells (Ousman and Kubes, 2012). As we and others have demonstrated that bOECs transplantation after SCI reduces inflammation into the spinal cord (Delarue et al., 2021; Khankan et al., 2016; Zhang et al., 2021), we hypothesize that it decreases systemic inflammation, limiting the disruption of BBB and brain reactivity. This hypothesis is indirectly confirmed by the decrease of hippocampal microglial reactivity observed in bOECs transplanted animals. Although most of the transplanted cells die within the first three weeks post-transplantation, it is possible that during this period, bOECs modulate levels of cytokines and growth factors in the brain, thereby reducing disruption to neurogenesis. This modulation is likely due to both a direct effect, through the release of anti-inflammatory cytokines and growth factors, and an indirect effect by stimulating spinal cord regeneration, which in turn reduces the release of pro-inflammatory cytokines and immune cell activation.

Finally, since physical activity stimulates learning and memory, we wondered whether these observations could be related to the functional recovery induced by the two types of cell transplantation. Although bOECs improved functional and tissue recovery after SCI, to our surprise, we did not observe any enhancement in motor abilities or spinal cord regeneration following MSCs transplantation (data not shown), which rules out this potential bias.

### Limitations of the study and Conclusion

To our knowledge, this is the first study to demonstrate that cells transplanted into the spinal cord can modulate the inflammatory response in the brain, thereby reversing the deleterious effects of injury on brain function following SCI. Furthermore, our study highlights this effect in two distinct transplantation paradigms, showing that these benefits occur both after the transplantation of differentiated cells, such as bOECs, and after the transplantation of stem cells, such as MSCs.

However, in our study, while we observed an immunomodulatory effect of these two cell types on microglial reactivity in the brain and on neuron density, we were unable to elucidate the precise mechanisms underlying these results. Specifically, it remains unclear whether the transplanted cells exert their effects via the bloodstream or the CSF, or which cytokines are modulated by these transplants. Therefore, further experiments are needed to precisely identify the mechanisms involved.

Nevertheless, our findings underscore the complexity of the processes that occur following SCI and cell transplantation. They also open the door to a new and yet unexplored field of research into the interactions between the brain and spinal cord under both physiological and pathological conditions.

## REFERENCES

Anjum, A., Yazid, M.D., Fauzi Daud, M., Idris, J., Ng, A.M.H., Selvi Naicker, A., Ismail, O.H.R., Athi Kumar, R.K., Lokanathan, Y., 2020. Spinal Cord Injury: Pathophysiology, Multimolecular Interactions, and Underlying Recovery Mechanisms. Int. J. Mol. Sci. 21, 7533. 10.3390/ijms21207533

Anthony, D.C., Couch, Y., (Prénom), 2014. The systemic response to CNS injury. Exp. Neurol. 258, 105–111. 10.1016/j.expneurol.2014.03.013

Assinck, P., Duncan, G.J., Hilton, B.J., Plemel, J.R., Tetzlaff, W., 2017. Cell transplantation therapy for spinal cord injury. Nat. Neurosci. 20, 637–647. 10.1038/nn.4541

Banks, W.A., Kastin, A.J., Gutierrez, E.G., 1994. Penetration of interleukin-6 across the murine blood-brain barrier. Neurosci. Lett. 179, 53–56. 10.1016/0304-3940(94)90933-4

Basu, J., Siegelbaum, S.A., 2015. The Corticohippocampal Circuit, Synaptic Plasticity, and Memory. Cold Spring Harb. Perspect. Biol. 7. 10.1101/cshperspect.a021733

Berdugo-Vega, G., Arias-Gil, G., López-Fernández, A., Artegiani, B., Wasielewska, J.M., Lee, C.-C., Lippert, M.T., Kempermann, G., Takagaki, K., Calegari, F., 2020. Increasing neurogenesis refines hippocampal activity rejuvenating navigational learning strategies and contextual memory throughout life. Nat. Commun. 11. 10.1038/s41467-019-14026-z

Capirossi, R., Piunti, B., Fernández, M., Maietti, E., Rucci, P., Negrini, S., Giovannini, T., Kiekens, C., Calzà, L., 2020. Early CSF Biomarkers and Late Functional Outcomes in Spinal Cord Injury. A Pilot Study. Int. J. Mol. Sci. 21. 10.3390/ijms21239037

Chalfouh, C., Guillou, C., Hardouin, J., Delarue, Q., Li, X., Duclos, C., Schapman, D., Marie, J.-P., Cosette, P., Guérout, N., 2020. The Regenerative Effect of Trans-spinal Magnetic Stimulation After Spinal Cord Injury: Mechanisms and Pathways Underlying the Effect. Neurother. J. Am. Soc. Exp. Neurother. 17, 2069–2088. 10.1007/s13311-020-00915-5

Chen, J., Wang, L., Liu, M., Gao, G., Zhao, W., Fu, Q., Wang, Y., 2022. Implantation of adipose-derived mesenchymal stem cell sheets promotes axonal regeneration and restores bladder function after spinal cord injury. Stem Cell Res. Ther. 13, 503. 10.1186/s13287-022-03188-1

Cherubini, E., Miles, R., 2015. The CA3 region of the hippocampus: how is it? What is it for? How does it do it? Front. Cell. Neurosci. 9. 10.3389/fncel.2015.00019

Craig, A., Guest, R., Tran, Y., Middleton, J., 2017. Cognitive Impairment and Mood States after Spinal Cord Injury. J. Neurotrauma 34, 1156–1163. 10.1089/neu.2016.4632

Delarue, Q., Mayeur, A., Chalfouh, C., Honoré, A., Duclos, C., Di Giovanni, M., Li, X., Salaun, M., Dampierre, J., Vaudry, D., Marie, J.-P., Guérout, N., 2019. Inhibition of ADAMTS-4 Expression in Olfactory Ensheathing Cells Enhances Recovery after Transplantation within Spinal Cord Injury. J. Neurotrauma 37, 507–516. 10.1089/neu.2019.6481

Delarue, Q., Robac, A., Massardier, R., Marie, J., Guérout, N., 2021. Comparison of the effects of two therapeutic strategies based on olfactory ensheathing cell transplantation and repetitive magnetic stimulation after spinal cord injury in female mice. J. Neurosci. Res. 99, 1835–1849. 10.1002/jnr.24836

Golestani, A., Shobeiri, P., Sadeghi-Naini, M., Jazayeri, S.B., Maroufi, S.F., Ghodsi, Z., Dabbagh Ohadi, M.A., Mohammadi, E., Rahimi-Movaghar, V., Ghodsi, S.M., 2022. Epidemiology of Traumatic Spinal Cord Injury in Developing Countries from 2009 to 2020: A Systematic Review and Meta-Analysis. Neuroepidemiology 56, 219–239. 10.1159/000524867

Gómez, R.M., Sánchez, M.Y., Portela-Lomba, M., Ghotme, K., Barreto, G.E., Sierra, J., Moreno-Flores, M.T., 2018. Cell therapy for spinal cord injury with olfactory ensheathing glia cells (OECs). Glia 66, 1267–1301. 10.1002/glia.23282

Guérout, N., Derambure, C., Drouot, L., Bon-Mardion, N., Duclos, C., Boyer, O., Marie, J.-P., 2010. Comparative gene expression profiling of olfactory ensheathing cells from olfactory bulb and olfactory mucosa. Glia 58, 1570–1580. 10.1002/glia.21030

Han, D., Wu, C., Xiong, Q., Zhou, L., Tian, Y., 2015. Anti-inflammatory Mechanism of Bone Marrow Mesenchymal Stem Cell Transplantation in Rat Model of Spinal Cord Injury. Cell Biochem. Biophys. 71, 1341–1347. 10.1007/s12013-014-0354-1

Khankan, R.R., Griffis, K.G., Haggerty-Skeans, J.R., Zhong, H., Roy, R.R., Edgerton, V.R., Phelps, P.E., 2016. Olfactory Ensheathing Cell Transplantation after a Complete Spinal Cord Transection Mediates Neuroprotective and Immunomodulatory Mechanisms to Facilitate Regeneration. J. Neurosci. Off. J. Soc. Neurosci. 36, 6269– 6286. 10.1523/JNEUROSCI.0085-16.2016

Li, Y., Ritzel, R.M., Khan, N., Cao, T., He, J., Lei, Z., Matyas, J.J., Sabirzhanov, B., Liu, S., Li, H., Stoica, B.A., Loane, D.J., Faden, A.I., Wu, J., 2020. Delayed microglial depletion after spinal cord injury reduces chronic inflammation and neurodegeneration in the brain and improves neurological recovery in male mice. Theranostics 10, 11376–11403. 10.7150/thno.49199

Liu, Y., Yang, X., He, Z., Li, J., Li, Y., Wu, Y., Manyande, A., Feng, M., Xiang, H., 2023. Spinal cord injury: global burden from 1990 to 2019 and projections up to 2030 using Bayesian age-period-cohort analysis. Front. Neurol. 14, 1304153. 10.3389/fneur.2023.1304153

Llorens-Martín, M., Jurado-Arjona, J., Avila, J., Hernández, F., 2015. Novel connection between newborn granule neurons and the hippocampal CA2 field. Exp. Neurol. 263, 285–292. 10.1016/j.expneurol.2014.10.021

Luna, A.C., Madeira, M.E., Conceição, T.O., Moreira, J.A., Laiso, R.A., Maria, D.A., 2014. Characterization of adipose-derived stem cells of anatomical region from mice. BMC Res. Notes 7. 10.1186/1756-0500-7-552

Maldonado-Bouchard, S., Peters, K., Woller, S.A., Madahian, B., Faghihi, U., Patel, S., Bake, S., Hook, M.A., 2016. Inflammation is increased with anxiety- and depression-like signs in a rat model of spinal cord injury. Brain. Behav. Immun. 51, 176–195. 10.1016/j.bbi.2015.08.009

McCandless, E.E., Piccio, L., Woerner, B.M., Schmidt, R.E., Rubin, J.B., Cross, A.H., Klein, R.S., 2008. Pathological Expression of CXCL12 at the Blood-Brain Barrier Correlates with Severity of Multiple Sclerosis. Am. J. Pathol. 172, 799–808. 10.2353/ajpath.2008.070918

Nakajima, H., Uchida, K., Guerrero, A.R., Watanabe, S., Sugita, D., Takeura, N., Yoshida, A., Long, G., Wright, K.T., Johnson, W.E.B., Baba, H., 2012. Transplantation of Mesenchymal Stem Cells Promotes an Alternative Pathway of Macrophage Activation and Functional Recovery after Spinal Cord Injury. J. Neurotrauma 29, 1614–1625. 10.1089/neu.2011.2109

Osburg, B., Peiser, C., Dömling, D., Schomburg, L., Ko, Y.T., Voigt, K., Bickel, U., 2002. Effect of endotoxin on expression of TNF receptors and transport of TNF-α at the blood-brain barrier of the rat. Am. J. Physiol.-Endocrinol. Metab. 283, E899–E908. 10.1152/ajpendo.00436.2001

Ousman, S.S., Kubes, P., 2012. Immune surveillance in the central nervous system. Nat. Neurosci. 15, 1096–1101. 10.1038/nn.3161

Prieto, G.A., Tong, L., Smith, E.D., Cotman, C.W., 2019. TNFα and IL-1β but not IL-18 suppresses hippocampal long-term potentiation directly at the synapse. Neurochem. Res. 44, 49–60. 10.1007/s11064-018-2517-8

Robac, A., Neveu, P., Hugede, A., Garrido, E., Nicol, L., Delarue, Q., Guérout, N., 2021. Repetitive Trans Spinal Magnetic Stimulation Improves Functional Recovery and Tissue Repair in Contusive and Penetrating Spinal Cord Injury Models in Rats. Biomedicines 9. 10.3390/biomedicines9121827

Tien, N.L.B., Hoa, N.D., Thanh, V.V., Thach, N.V., Ngoc, V.T.N., Dinh, T.C., Phuong, T.N.T., Toi, P.L., Chu, D.T., 2019. Autologous Transplantation of Adipose-Derived Stem Cells to Treat Acute Spinal Cord Injury: Evaluation of Clinical Signs, Mental Signs, and Quality of Life. Open Access Maced. J. Med. Sci. 7, 4399–4405. 10.3889/oamjms.2019.843

Wagner, F.B., Mignardot, J.-B., Le Goff-Mignardot, C.G., Demesmaeker, R., Komi, S., Capogrosso, M., Rowald, A., Seáñez, I., Caban, M., Pirondini, E., Vat, M., McCracken, L.A., Heimgartner, R., Fodor, I., Watrin, A., Seguin, P., Paoles, E., Van Den Keybus, K., Eberle, G., Schurch, B., Pralong, E., Becce, F., Prior, J., Buse, N., Buschman, R., Neufeld, E., Kuster, N., Carda, S., von Zitzewitz, J., Delattre, V., Denison, T., Lambert, H., Minassian, K., Bloch, J., Courtine, G., 2018. Targeted neurotechnology restores walking in humans with spinal cord injury. Nature 563, 65–71. 10.1038/s41586-018-0649-2

Wang, X., Jiang, C., Zhang, Yongyuan, Chen, Z., Fan, H., Zhang, Yuyang, Wang, Z., Tian, F., Li, J., Yang, H., Hao, D., 2022. The promoting effects of activated olfactory ensheathing cells on angiogenesis after spinal cord injury through the PI3K/Akt pathway. Cell Biosci. 12, 23. 10.1186/s13578-022-00765-y

Whitlock, J.R., Heynen, A.J., Shuler, M.G., Bear, M.F., 2006. Learning Induces Long-Term Potentiation in the Hippocampus. Science. 10.1126/science.1128134

Zhang, L., Zhuang, X., Kotitalo, P., Keller, T., Krzyczmonik, A., Haaparanta-Solin, M., Solin, O., Forsback, S., Grönroos, T.J., Han, C., López-Picón, F.R., Xia, H., 2021. Intravenous transplantation of olfactory ensheathing cells reduces neuroinflammation after spinal cord injury via interleukin-1 receptor antagonist. Theranostics 11, 1147– 1161. 10.7150/thno.52197

Zhao, M., Choi, Y.-S., Obrietan, K., Dudek, S.M., 2007. Synaptic Plasticity (and the Lack Thereof) in Hippocampal CA2 Neurons. J. Neurosci. 27, 12025–12032. 10.1523/JNEUROSCI.4094-07.2007

